# Axon Guidance Signaling Modulates Epithelial to Mesenchymal Transition in Stem Cell-Derived Retinal Pigment Epithelium

**DOI:** 10.1101/264705

**Authors:** Srinivas R. Sripathi, Melissa M. Liu, Ming-Wen Hu, Jun Wan, Jie Cheng, Yukan Duan, Joseph Mertz, Karl Wahlin, Julien Maruotti, Cynthia A Berlinicke, Jiang Qian, Donald J. Zack

**Affiliations:** Department of Ophthalmology, Wilmer Eye Institute, The Johns Hopkins University School of Medicine Baltimore, MD 21205, USA.; Department of Medical and Molecular Genetics, Indiana University School of Medicine, Indianapolis, IN 46202, USA; Shiley Eye Institute, University of California, San Diego, LA Jolla, CA 92093, USA.; Phenocell, Evry cedex, France; Solomon H. Snyder Department of Neuroscience, Department of Molecular Biology and Genetics, Institute of Genetic Medicine, The Johns Hopkins University School of Medicine, Baltimore, MD 21205, USA.

## Abstract

The critical role of epithelial to mesenchymal transition (EMT) in embryonic development, malignant transformation, and tumor progression has been well studied in normal and cancerous tissues and cells. Interestingly, EMT has also been reported to play a key role in the early progression of several retinal degenerative diseases, including scarring associated proliferative vitro-retinopathy (PVR), choroidal neo-vascularization induced “wet” age-related macular degeneration (AMD) and diabetic retinopathy (DR). Despite these studies, many questions remain unexplored regarding EMT-associated retinal pigment epithelium (RPE) degeneration and dysfunction. We hypothesize that RPE cells undergo EMT prior to cell death during the progression of atrophic “dry” AMD. Utilizing human stem cell-derived RPE (hRPE) as a model to study RPE EMT, we optimized two independent but complementary RPE EMT induction systems: 1) enzymatic dissociation of hRPE monolayer cultures and 2) co-treatment of hRPE monolayer cultures with transforming growth factor beta (TGF-β) and the inflammatory cytokine, tumor necrosis factor alpha (TNF-α). To further understand the molecular mechanisms of RPE EMT regulation, we performed an RNA-Sequencing (RNA-Seq) time course examination across 48 hours beginning with EMT induction. Our transcriptome profiling provides a comprehensive quantification of dynamic signaling events and associated biological pathways underlying RPE EMT and reveals an intriguing significance for widespread dysregulation of multiple axon guidance molecules in this process.

## INTRODUCTION

Epithelial to mesenchymal transition (EMT), is defined as the repression of epithelial characteristics and acquisition of a mesenchymal phenotype. It is well known that different transcription factors tightly regulate EMT through a variety of signaling pathways^1-3^. During early progression of EMT, before complete transformation to a mesenchymal phenotype, epithelial cells re-organize their cytoskeletal architecture resulting in loss of integral tight junctions, and exhibit disrupted apicobasal polarity and changed cell morphology through altered gene expression and a subsequent protein level change ^4-5^. Previously, EMT had only been observed during embryonic development, tissue fibrosis, tumor dissemination and most commonly, metastasis, in which epithelial cells lose cadherin and integrin to acquire a motile phenotype and invade neighboring healthy cells^6-8^. Besides the crucial role of EMT during development and tumor progression, this transition has recently been reported to be involved in retinal degenerative diseases including fibrotic scarring associated proliferative vitro-retinopathy (PVR)^9-11^, choroidal neovascularization (CNV) associated wet or vascular age-related macular degeneration (AMD)^12-14^ and the progression of dry or atrophic AMD^15^. In each of these diseases, EMT has been associated with the dysfunction of retinal pigment epithelium (RPE). RPE is a highly pigmented, hexagonally packed cuboidal monolayer of cells, which resides on Bruch's membrane between the neural retina and choriocapillaris. Healthy RPE cells exhibit apicobasal polarity and tight connections with photoreceptor cells of the retina, for which they perform crucial support functions, including phagocytosis of rod outer segments, synthesis of vitamin-A metabolites (retinoid recycling) and protection against reactive oxygen species^16-17^. During EMT progression, RPE loses tight junction integrity and pigmentation and acquires a fibroblast-like phenotype. The resulting collective loss of function leads to early events associated with RPE degenerative diseases, and more specifically AMD^15^, in which damaged RPE cells can no longer properly maintain the health and function of neighboring photoreceptor cells. Therefore considerable attention has been devoted to studying RPE cell biology and pathology in AMD. Past studies were limited by their use of animal models, human fetal samples, cadavers, and donor specimens. Efficient methods have recently been developed for differentiating human ES and iPS cells into “RPE” (hRPE)^18-19^ that closely mimics native RPE cells at morphological, biochemical, molecular and functional levels. These methods have provided a novel model system in which to examine the key factors involved during both stages of differentiation and de-differentiation as they occur in development and disease progression. To better understand the complexity of EMT in RPE, to it is important to address crucial questions such as which early regulatory factors and key signaling pathways are candidates for further inhibition of EMT progression. EMT can be induced and regulated by various growth factors and cytokines, TGF-β signaling specifically plays a critical role in embryonic development, tumor progression and tissue fibrosis^20-21^. A resemblance to the EMT that occurs during tumor progression is seen in our selective gene expression data on enzymatic dissociation, TGF-β/ TNF-α induced RPE EMT.

Our preliminary studies show that several mesenchymal and epithelial-specific transcription factors (TF) are differentially regulated during the EMT progression in iPS-derived RPE cells. After confirming that all the RPE-specific genes are significantly downregulated to basal level during EMT progression, we hypothesized that through study of the RPE EMT transcriptome would not only provide a foundation for understanding the RPE trans-differentiation process but also improve the potential of RPE transplantation therapeutics and drug discovery approaches to halt and even revert EMT in retinal degenerative diseases. To investigate this, we used deep RNA-Sequencing (RNA-Seq) transcriptomic profiling as an advanced strategy for determining and understanding the various signaling events influencing RPE phenotypic paradigm. Recently, RNA-Seq has been widely used to trace the whole transcriptome in a large variety of cells and tissues of disease models at an unforeseen depth and sensitivity. The system wide molecular signature of RPE EMT is still uncharted territory. To date, there has been no in-depth temporal transcriptomic comparison between normal and EMT-acquired RPE cells derived from human stem cells. Here we demonstrate that integrative proteo-genomic profiles from stem cell-derived RPE monolayers compared to those with EMT induced by enzymatic dissociation and altered TGF-β signaling adds comprehensive information to evaluate the progression of RPE EMT in vitro. Altogether, our transcriptomic data provide a solid platform for predicting the most functionally altered biological pathways and factors relevant to early RPE EMT initiation during the progression of the retinal degenerative disease, specifically AMD.

## Results

### hiPS RPE differentiation, enzymatic dissociation, and TGF-β signaling induced RPE EMT

hiPS cells were differentiated into mature RPE monolayers using our previously published methods^18-19^. During differentiation, the morphology of hiPSCs after 50 days in differentiation medium shows pigmented colonies (Fig 1A). Flow cytometry shows that more than 90 % expression of the pigment-related melanocyte-specific protein (PMEL17) and the RPE-specific key visual cycle protein (RPE65) from differentiating RPE cells (Fig 1B-C). To explore dedifferentiation of hiPS-RPE monolayers, we induced EMT by detaching the RPE monolayer cultures from the culture substrate and dissociated them to single cells by proteo-collagenolytic enzymes (AccuMAX). We observed EMT related phenotypic changes at 3-48 hrs time point (Fig 1D). We also induced EMT by co-treatment with TGF-β/TNF-α and AMD-associated oxidative stressor. We repeatedly observed the elevation of key EMT-related transcription factors such as Snail family zinc finger (SNAI1, SNAI2), Zinc-finger E-box-binding (ZEB), basic-loop-helix transcription 1 (TWIST1) and transcription factor-3 (TCF3) (Fig 1). We also observed significant upregulation of downstream EMT-associated genes including α-smooth muscle actin (ACTA2), vimentin (VIM), fibronectin (FN1) and N-Cadherin (CDH2). Conversely, RPE-specific factors such as pre-melanosome protein (PMEL17), microphthalmia-associated transcription factor (MITF), chloride channel-related protein bestrophin 1 (BEST1), tyrosinase (TYR), retinaldehyde-binding protein 1 (RLBP1) and retinal pigment epithelium-specific protein of 65 kDa (RPE65) were dramatically down-regulated. Further, we identified elevated levels of transcripts from other key EMT pathways such as NOTCH1, HES1, HEY1 (notch signaling) TGFβ1, NFkβ1, HMGA2, GSk3B, β-catenin, glioma1 (TGFβ, WNT, NFkβ signaling) during the initiation and progression of RPE EMT (SI Fig. 1). Post-enzymatic treatment of RPE monolayers followed by re-plating the cells showed that the cells lose their RPE-like characteristics and exhibit an altered phenotype including loss of pigmentation and acquisition of elongated fibroblast morphology and migratory behaviors (Fig 1 E-G). Consistent with the implication of EMT playing a role RPE dysfunction during early progression of AMD^15^, we also examined the expression of key EMT and RPE factors after treatment of RPE monolayers with AMD-associated oxidative stressors such as paraquat (PQ), tert-butyl hydroperoxide (tBuOOH) and cigarette smoke extract (CSE) (data not shown).

**Figure 1.**
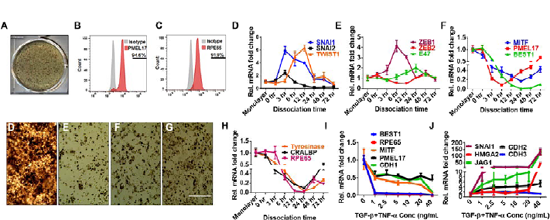
hiPS RPE differentiation, characterization and EMT induction. A) Morphology of hiPSC after 45d of differentiation. B) Flow-cytometric analysis of RPE65 and PMEL-17 expression from 2-month-old RPE. C) Bright field images of the RPE monolayer and after enzymatic dissociation to induce EMT. D) Differential expression of key EMT and RPE factors were measured by qRT PCR analysis.

### Distinct transcriptomes of enzymatic dissociation and TGF-β signaling induced RPE EMT

For a comprehensive examination of gene expression changes that occur during RPE EMT, we performed deep paired-end RNA-Seq on dissociated hRPE monolayer cultures over time (3hr, 12hr and 48 hr) and on hRPE monolayer cultures co-treated with different doses of TGF-β/TNF-α(1-40 ng/mL). We obtained an average of 13.4 million reads per sample (dissociation) with the average mapping rate of 64.2 % to the human genome (NCBI build37.2) and an average of 15 million reads per sample (TGF-β/TNF-α) with an average mapping rate of 53.11%. Unsupervised hierarchical clustering showed that the untreated and EMT induced populations were grouped together and well separated (Fig 1 B-C). Differentially expressed genes (DEGs) were evaluated by applying a stringent statistic threshold of a greater than or equal to two-fold change (FC), a false discovery rate (FDR) of <0.05 and a p-value of <0.05 (student t-test). As a result, we identified a list of 5397, and 1439 DEGs from enzymatic dissociation and TGF-β/TNF-α induced EMT (Fig 1 B-C) respectively. We identified a consensus of 780 genes showing significant overlap between dissociation and TGF-β/TNF-α induced EMT. Differentially regulated top 40 genes from both EMT types were shown in heatmaps (Fig 2 E-J)To further discern the biological changes, altered canonical pathways, upstream transcriptional regulators and gene networks that regulate RPE EMT, we performed Ingenuity Pathway Analysis (IPA) and gene ontology (GO) analysis. IPA identified several pathways significantly altered by both EMT induction methods (Fig 3 A-C). The list of top canonical pathways were calculated using Benjamini & Hochberg (BH) adjusted p-value, and the number of identified molecules were shown from each RPE EMT category, and that overlapped in both EMT types. Additionally, to investigate the cascades of transcription regulators that link DEGs and their involvement in various signaling events during RPE EMT, we performed upstream regulator analysis (URA), which computes potential upstream transcriptional regulators that could have brought about the observed patterns, based on prior information. The upstream transcription algorithm uses statistical methods such as overlap p-value and an activation z-score to predict the key upstream regulators that alters gene expression. These regulators include “transcription regulators,” which are predominantly DNA-associated transcription factors, various kinases, miRNAs, translational regulators, growth factors, cytokines, and chemical drugs including numerous kinase inhibitors that have been observed experimentally to alter gene expression^22-23^. We clustered the resulting transcription regulators based on activation Z-score and p-value overlap.

**Figure 2.**
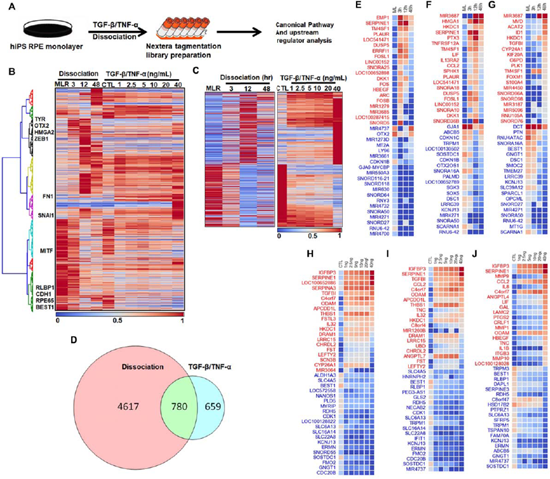
Temporal transcriptomic profiling of enzymatic dissociation induced RPE EMT. A. Schematic work flow for RNA-Seq analysis. B. Hierarchical clustering of log2-transformed ratios. Average abundances of differentially expressed genes showing significant differences across time point after dissociation and TGF-β/TNF-α induced EMT in hiPS RPE. C. Average abundances of differentially expressed overlapped genes from dissociation and TGF-β/TNF-α induced EMT in hiPS RPE. D. Venn diagram represents the differentially regulated genes that share the commonality between dissociation and TGF-β/TNF-α induced EMT in RPE. E-G. Heatmap of the top 40 differentially regulated RPE EMT genes from RNA-Seq analysis

**Figure 3.**
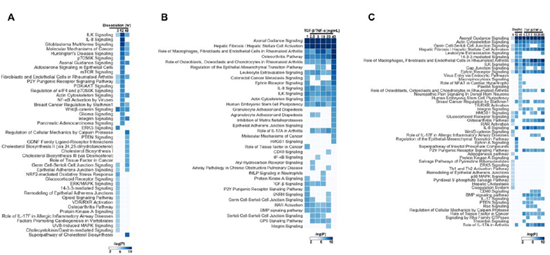
IPA analysis of top canonical pathways altered in dissociation and TGF-β/TNF-α induced RPE EMT. Top cellular pathways were predicted based on the regulation of highly enriched genes that changed in abundance (activated or inhibited) from undissociated monolayers to dissociated (3 hr, 12 hr and 48 hr) cells (A), TGF-β/TNF-α treated RPE monolayers (B) and overlapped from both EMT types (C). −log 10 overlapping p values were computed using Fisher’s exact test and enriched canonical pathways were identified based on the observed number of genes for each pathway, relative to the number expected by chance.

### Axon guidance signaling modulates RPE EMT in stem cell-derived RPE monolayers

Gene Ontology (GO), Ingenuity Pathway Analysis (IPA) and Kyoto Encyclopedia of Genes and Genomes (KEGG) pathway enrichment analysis revealed that the axon guidance signaling pathway is one of the most highly enriched pathways from our RPE EMT dataset with a −log(p-value) between 6.4-14.8. A axon guidance signaling molecules have previously been are implicated in the progression of EMT during tumor metastasis.^24-26^ Actin cytoskeleton, Interleukin, ephrin receptor signaling pathways were also enriched. Based on IPA and GSEA findings indicating very significant altered expression of axon guidance-related genes, we analyzed RNA-Seq data for coordination of major guidance cue ligand/receptor pairings, such as 1) semaphorins, which bind with plexins/neuropilins receptors, 2) netrins, which bind with DCC *(Deleted in* Colorectal Cancer)/UNC5 (Un-Coordinate loss-of-function) receptors, 3) slits, which bind with ROBO (Roundabout) receptors, and 4) ephrins, which bind with Eph receptors.^27-28^ Interestingly, widespread dysregulation of multiple axon guidance ligand-receptor pairs was observed, with pairs such as SEMA4B-PLXNB1, SEMA4F-NRP2, NTN4-UNC5B, SLIT3-ROBO4, EFNA5-EPHA5, EFNB1-EPHB1, and EFNB2-EPHB2 exhibiting significant upregulation, while NTNG1-UNC5D and SLIT2-ROBO1 showing down-regulation from dissociation induced EMT. SEMA3E-PLXND1, SEMA6A-PLXNA2, EFNB1-EPHB1, and EFNB2-EPHB2 pairs were up-regulated, and SEMA4A-PLXNB1, SEMA4E, PLXNB1,SEMA4G-PLXNB2, NTNG1-UNC5D, SLIT1-ROBO2, and EFNA5-EPHA5 were down-regulated in TGF-p/TNF-a induced EMT. To further validate the differential expression of axon guidance molecules measured in RNA-Seq, we quantified expression of the implicated axon guidance molecules using qRT-PCR (Fig 5 C-O), and found that genes encoding Semaphorin family factors (SEMA3A, SEMA3D, SEMA4A, SEMA6D), canonically believed to be involved in the inhibitory signaling of axon growth cones, were up-regulated., Conversly, a axon guidance attractive receptor molecules (EPHB1, EPHB2) which interact with ephrins (EFNB1,EFNB2), were down-regulated in RPE EMT.

### IPA upstream regulator analysis identifies small molecule inhibitors that restore key RPE genes by inhibiting RPE EMT

To validate the top hits from the chemical kinase inhibitor list generated by the upstream regulator analysis (Fig. 6), we treated hRPE monolayers with the identified small molecules prior to EMT induction by enzymatic dissociation or TGF-β/TNF-α treatment and analyzed them qPCR. Briefly, RPE monolayers were treated in triplicate with a broad dose range (80 μM, 40 μM, 20 μM, 10 μM, 5 μM, 2.5 μM, 0.6 μM, 0.05 μM) of kinase inhibitors including PD98059 (MAPK (Erk) kinases MEK1), LY294002 (PI3K inhibitor), U0126 (MEK1/MEK2 inhibitor), SB203580 (p38 MAPK inhibitor) and Tyrphostin AG1478 (EGFR inhibitor) followed by the induction of EMT. Interestingly, our qPCR analysis shows that U0126, which inhibits the activation of MAPK (ERK 1/2) by inhibiting the kinase activity of MAP Kinase Kinase (MAPKK or MEK 1/2), potently restores the expression of key RPE factors such as BEST1 (36fold), RPE65 (18-fold) and LRAT (6-fold) by regulating EMT factors CDH2 (15-fold decreased) and CDH1 (5-fold increased). Collectively, these data suggest that blockage of MEK1/2 signaling may be a potent pharmacological target for inhibition of EMT in RPE.

## Discussion

In the current study, we analyzed temporal, dose-dependent transcriptome-wide changes in gene expression abundance in response to enzymatic dissociation and TGF-β/TNF-α stimulation of hiPS-RPE cells. Our data show that the expression of key functional RPE genes such as MITF, PMEL17, BEST1 and RPE65, were significantly down-regulated from both dissociation and TGF-β signaling induced RPE EMT, however the differential regulation of key EMT factors are not common from each EMT type. For instance, known EMT factors that are elevated during malignant metastasis such as SNAI2, TWIST1, ZEB1 significantly alter during dissociation induced EMT but not from TGF-β/TNF-α induced EMT. Our RNA-Seq analysis identifies many previously unreported key factors that regulate early and late onset of RPE EMT; We validated several genes by qRT PCR analysis, and also we provided the evidence that strong correlation of these selected genes with transcriptomic data. Thus, our data are consistent with previously reported EMT progression during tumor metastasis, and this study might provide a valuable resource for exploring the pathological role od RPE EMT during the early progression of serval RPE involved retinal degenerative diseases.

Although our results show the broad range of signaling pathways contributing to the induction of EMT in RPE, each pathway might distinctly regulate downstream RPE gene expression. Investigation of altered canonical pathways, upstream transcription regulators and network analysis data suggested that several transcription factors and other cofactors such as kinases, growth factors, cytokines, enzymes, miRNAs, could contribute to the dissociation and TGF-β signaling induced RPE EMT gene expression changes (Fig. 4). IPA based canonical pathway analysis shows that integrin-linked kinase (ILK) signaling, which mediates several key events including cell survival, proliferation and differentiation is top to enriched from dissociation induced EMT. ILK signaling regulates the cross-talk between E-cadherin and integrin^3^, and the increased expression of ILK results in the down-regulation of E-cadherin through the activation of β-catenin and nuclear factor (NF)-kB^29-31^. We observed that dissociation and TGF-β/TNF-α induced RPE EMT increased the abundances of E-cadherin, β-catenin, and NF-kB1 (SI Fig 1). Transcription factors such as NFKB1, STAT1, STAT3, RELA, JUN, SP1, HIF1A, NUPR1 were activated, and OTX2, MEOX2, KLF2, BCL6, ZFP36, TAF4 were inhibited throughout the dissociation time course, and TGF-β/TNF-α dose response as identified by IPA generated upstream regulator analysis (Fig 4). Additionally, IPA detected numerous kinases, that might play key role towards initiating RPE EMT, including INSR, IPMK, (dissociation); PRKCD, PRKCE, MAPK3, IKBKB (TGF-β/TNF-α); SRC, RAF1, MAP3K1, MAP3K4, MAP3K14, FGFR1, ERBB2, EGFR, MKNK1 (from both dissociation and TGF-β/TNF-α induced EMT). Further, upstream regulator analysis also predicts several growth factors (EGF, FGF2, AGT, HGF, VEGFA, NRG1, PDGFB, NRG1, NOG, DKK1, WISP2); cytokines (CSF1-2, CCL5, SPP1, TSLP, TNF, CD40LG, OSM, IL1-6,13, 24, IFNG, EDN1); and enzymes (RHOA, KRAS, TGM2, IRS1, NCF1, TRAF, HRAS, PTGS2, FN1, TET2, SNCA, PIN1), that could regulate early RPE EMT. Recent studies reveal that RPE EMT is also posttranscriptionally regulated by multiple non-coding miRNAs^32^'^35^. Although our RNA-Seq analysis identified only the differential regulation of miRNA host genes during RPE EMT, IPA upstream regulator analysis suggested that several miRNAs could contribute to the changes in RPE EMT gene expression. mir-210, mir-26, mir-17-5p, mir-141-3p, mir-30c-5p were activated, but mir-122, mir-146, miR-16-5p, miR-1-3p, miR-124-3p, miR-30c-5p, let-7a-5p were inhibited through time and dose-dependent (Fig 4). Interestingly, it’s been well reported that some miRNAs from the above list alter EMT signaling by targeting various factors during various cancer metastasis. For instance, miR-210 (ovarian cancer^36^), miR-122 (hepatocellular carcinoma^37^), miR146 (non-small cell lung cancer^38^), miR-124-3p (bladder cancer^39^), miR-30c-5p (gastric cancer^40^) modulates EMT during tumor invasion and metastasis.

**Figure 4.**
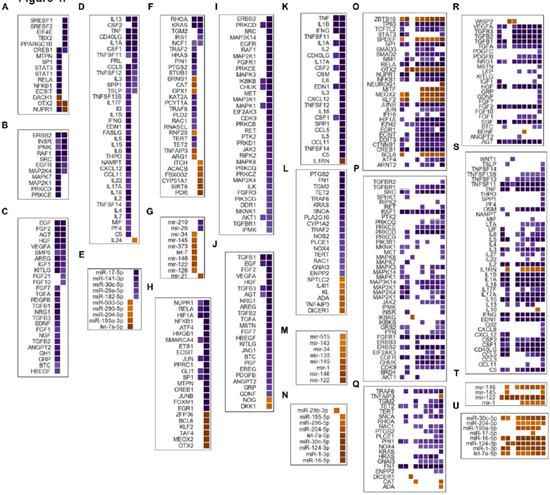
Upstream regulator analysis dissociation and TGF-β/TNF-α induced RPE EMT. Heatmap of the activation scores for upstream transcriptional regulators predicted to be activated (violet) or inhibited (yellow) after enzymatic dissociation in at least on time point. IPA uses activation Z-score as a statistical measure of the match between expected relationship direction and observed changes in gene expression. Z-score of >2 or <-2 is considered as a significant. regulator such as different transcription regulators, kinases, growth factors, cytokines, enzymes, miRNAs, predicted kinase inhibitors etc. Only genes with statistically significant changes at a FDR of 5% (q<0.05) were included in the analysis.

**Figure 5.**
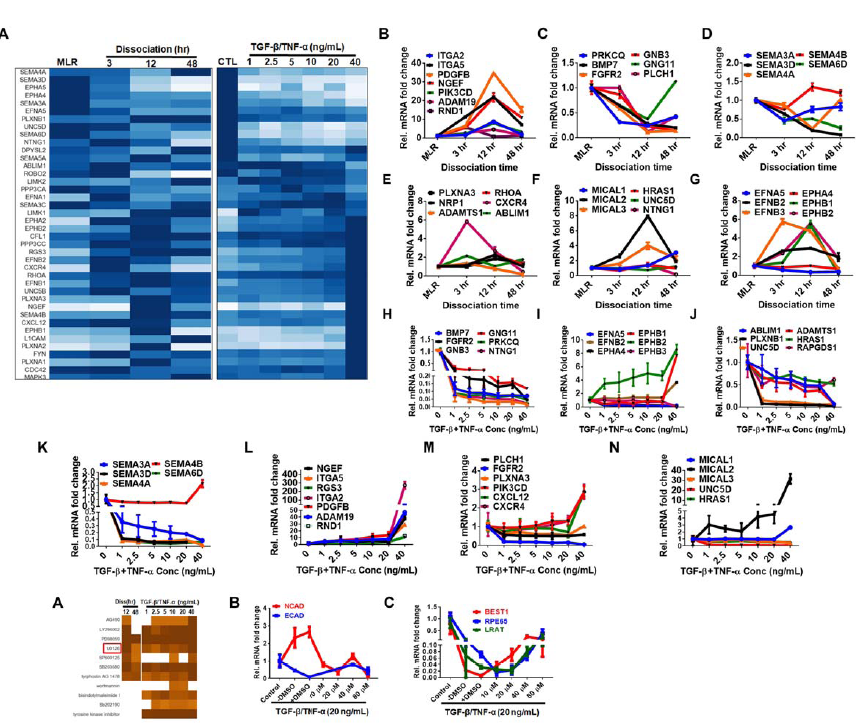
Enrichment of axon guidance genes from dissociation and TGF-β/TNF-α induced RPE EMT. A. Heatmap of multiple axon guidance molecules identified by KEGG and IPA. B. Validation of axon guidance molecules by qRT-PCR analysis (C-P).

The involvement of axon guidance molecules modulating EMT during embryonic development, tumor invasion, and metastasis remains unexplored. Here we demonstrated a strong and unexpected association between axonal guidance signaling and RPE EMT. Our transcriptome analysis shows the valid cross talk by the same ligand-receptor interaction between the key axonal guidance molecules. For instance, SEMA4B, SEMA4F SEMA3E, SEMA6A and their respective binding receptors PLXNB1, NRP2, PLXND1, PLXNA2 were significantly up-regulated during RPE EMT. Precisely, SEMA3E/PLXND1signaling modulated endothelial cell migration involved in vessel formation in cardiovascular models^41^, development of axon tracts in the forebrain^42^ and synaptic specificity in sensory-motor connections^43^ is reported. Our data shows that SEMA4A/4E/4G and its possible binding receptors PLXNB1/B2 were down-regulated during TGF-β/TNF-α induced EMT. The binding of SEMA4A to B-type plexins from our data is consistent with the recent studies that showed the binding of Sema4A with PLXND1 suppresses the VEGF mediated endothelial cell proliferation and migration^44^, activates growth cone collapse through the inhibition of R-Ras activity in mouse hippocampal neurons^45^. Further, down-regulation of SLIT1-ROBO2 interaction in RPE EMT supports the evidence that Slit family regulated intra-retinal axon outgrowth through ROBO2 towards mediating the polarity of retinal ganglion cell (RGC) ^43^. Interestingly, ephrin ligand, ephrin-A5 (EFNA5) and its receptor EPHA5 show the decreased expression form both dissociation and TGF-β/TNF-α induced EMT, which correlates with the involvement of ephrin signaling with the maintenance of cell morphology and adhesion^46^. Furthermore, our IP A upstream regulator analysis identifies the small molecule kinase inhibitor (U0126) partially reverses TGF-β/ TNF-α induced RPE EMT (Fig 6). TGF-β signaling induces the phosphorylated Erk and Erk kinases activity, and it is shown that U0126 blocks the activated Erk in NMuMG cells in vitro and further inhibited EMT^47^. Overall, our unsupervised transcriptome analysis of EMT in RPE unveils different early and late intrinsic molecular pathways and their link to the biological process of RPE EMT.

## METHODS

### Human Pluripotent Stem Cell (hPSC) Culture and Differentiation into RPE

Human hPSC lines were cultured and differentiated into RPE as previously described^18-19^. Briefly, the hPSC line EP1 were maintained on growth factor-reduced Matrigel (BD Biosciences) in mTeSR1 medium (Stem Cell Technologies), in a 10% CO_2_ and 5% O_2_ incubator and amplified by clonal propagation using ROCK pathway inhibitor blebbistatin (Sigma). For differentiation, hPSCs were plated at higher density (25,000 cells per cm^2^) and maintained in mTeSR1 to form a monolayer and the culture medium was replaced with differentiation medium (DM) for 40-45 days.

### Flow cytometry analysis

Immunostaining for RPE-specific markers were performed using the IntraPrep Permeabilization kit (Beckman Coulter) as per the manufacturer’s instructions. Primary antibody concentration was one μg per 1 million cells for mouse anti-PMEL17 (Abcam), mouse anti-RPE65 (Abcam). Goat anti-mouse conjugated to Alexa 647 (Invitrogen) was used as a secondary antibody. Nonspecific, species-appropriate isotype control was included in all flow cytometry experiments and stained cells were analyzed using a C6 flow cytometer (Accuri). Further histogram analyses were performed using FloJo software (FloJo, Ashland, OR).

### RNA isolation and quantitative RT-PCR

Total RNA from RPE cells were extracted using Isolate II RNA mini kit (Bioline) and reverse-transcribed (High Capacity cDNA kit; Applied Biosystem) before qPCRs were performed with so advanced master mix (Bio-Rad). Quantitative PCR samples were run in biological triplicates and expression levels normalized using the geometric mean of reference genes including GAPDH, FBXL12, SRP72, and CREBBP. Gene-specific primers sequences were included in supplementary Table 1.

**RNA-Seq and Data Processing.** First strand cDNA synthesis was performed with 195 ng total RNA using anchored oligo-dT and SuperScript III First-Strand Synthesis SuperMix (ThermoFisher, Waltham, MA, USA). Second strand cDNA synthesis was performed using RNase H, DNA Polymerase I, and Invitrogen Second Strand Buffer (ThermoFisher, Waltham, MA, USA). Double-stranded cDNA was purified using DNA Clean & Concentrator-5 (Zymo Research, Irvine, CA, USA). Library preparation was performed using the Nextera XT DNA Library Preparation Kit (Illumina, San Diego, CA, USA). Libraries were cleaned using Agencourt AMPure XP beads according to manufacturer’s instructions (Beckman Coulter, Brea, CA, USA). Libraries were evaluated by the High Sensitivity DNA Kit on the 2100 Bioanalyzer. They were then multiplexed and sequenced on an Illumina HiSeq with 50 bp paired-end reads. Reads were aligned to NCBI build 37.2 using Tophat (v2.1.0). Cuffquant and Cuffnorm (Cufflinks v2.2.1) were used to quantify expression levels and calculate normalized FPKM values^49^.

### Small-Molecule treatment

Chemical kinase inhibitors (Sellectchem) were reconstituted in DMSO (10 mM and 200 μM stocks) and loaded into Echo qualified 384-well polypropylene microplates and dispensed precisely in multiples of 2.5 nL droplets at the desired concentrations using acoustic liquid handling system ECHO 550 (Labcyte, CA).

### RNA-Seq data analysis

We perfoemed student t-test to find differentially expressed genes. For enzymatic dissociation data, the thresholds were log2 fold change > 1 and FDR <0.3. For TGF-β/TNF-α induced EMT data, the thresholds were log2 fold change > 1 and FDR <0.1. For unsupervised hierarchical clustering, pearson’s correlation coefficient was used to construct the linkage matrix, and used ward method for calculating distance between clusters.

**Figure 6. Small molecule treatment restores key RPE factors by inhibiting RPE EMT.** A. Upstream regulator analysis shows the heatmap of the activation Z-scores for kinase inhibitors predicted to be inhibited (brown) or activated (blue) by dissociation (3,12 hr) or stimulus with TGF-β/TNF-α. (1-40 ng/mL) B-C. Restoration of RPE/EMT markers with the treatment of U0126 is analyzed by qRT PCR.

## Acknowledgements

The authors thank Noriko Esumi, Xue Yang, Xiaomei Han, Xitiz Chamling, John Fuller, Pingwu Zhang, Bibhudatta Mishra, Claire Wegner for their insightful suggestions and discussions. This work is supported by grants from NIH, Maryland Stem Cell Research Foundation, Beckman Foundation, Foundation Fighting Blindness, Research to Prevent Blindness, Bright Focus Foundation and generous gifts from the Guerrieri Family Foundation and Mr. and Mrs. Robert and Clarice Smith, and the Raab Family Foundation.

## Author Contributions

D.J.Z., S.R.S designed the research. S.R.S., M.L., J.C., performed the experiments. D.J.Z., J.Q., S.R.S., M.M.L., M.H., WJ., J.C., J.L.M., C.A.B., analyzed the data. K.J.W., J.M., contributed samples, new reagents/analytic tools. S.R.S and D.J.Z., wrote the paper.

